# Cell based strain stiffening of a non-fibrous matrix as organizing principle for morphogenesis

**DOI:** 10.1101/816496

**Authors:** Daniel Rüdiger, Kerstin Kick, Andriy Goychuk, Angelika M. Vollmar, Erwin Frey, Stefan Zahler

**Affiliations:** Department of Pharmacy, Pharmaceutical Biology, Ludwig-Maximilians-Universität München, Butenandtstraße 5-13, 81377 Munich, Germany; Arnold Sommerfeld Center for Theoretical Physics and Center for NanoScience, Department of Physics, Ludwig-Maximilians-Universität München, Theresienstraße 37, 80333 Munich, Germany

## Abstract

Endothelial tube formation on a reconstituted extracellular matrix (Matrigel) is a well-established *in vitro* model for studying the processes of angiogenesis and vasculogenesis. However, to date, the organizing principles that underlie the morphogenesis of this network, and that shape the initial process of cell-cell finding remain elusive. Furthermore, it is unclear how *in vitro* results extrapolate to *in vivo* morphogenesis. Here, we identify a mechanism that allows cells to form networks by mechanically reorganizing and stiffening their extracellular matrix, independent of chemical guidance cues. Interestingly, we find that this cellular self-organization strongly depends on the connectivity and topology of the surrounding matrix, as well as on cell contractility and cell density. Cells rearrange the matrix, and form bridges of matrix material that are stiffer than their surroundings, thus creating a durotactic track for the initiation of cell-cell contacts. This contractility-based communication via strain stiffening and matrix rearrangement might be a general organizing principle during tissue development or regeneration.

**Significance Statement:** In addition to chemotactic gradients, biomechanical cues are important for guiding biological pattern formation. Self-assembly of cells has often been ascribed to reorganization of collagen fibres in the extracellular matrix. However, the basement membrane surrounding vascular cells, is *per se* non-fibrous. Here, we find that this difference in matrix topology can crucially influence cell behaviour and pattern formation. In a homogeneously elastic environment like the basement membrane, endothelial cells rearrange extracellular matrix proteins by contractile force, forming stiff intercellular bridges as tracks for cell-cell contacts. Our findings shine some light why there is a lot of merit in having multiple approaches to matrix elasticity (like continuum theories or dilated network approaches). Our observations might help to understand why vascular nets look different in different tissues and after rearrangement of the extracellular matrix during disease.

## Introduction

Angiogenesis, the sprouting of new vessels from pre-existing ones, and vasculogenesis, the *de novo* formation of vessels from blood islands during embryogenesis are crucial for any tissue growth – be it physiological (during development or wound healing) or pathophysiological (e.g. in solid tumours). Since the discovery of Vascular Endothelial Growth Factor (VEGF), the concept that vascular networks are shaped and driven by chemotactic gradients became predominant (1). The clinical success of anti-VEGF therapy further fuelled this hypothesis (1).

In recent years it became more and more clear that biophysical cues, and especially interactions of cells with the extracellular matrix (ECM), are important determinants for morphogenesis of tissues in health and disease (2, 3), especially in organs showing tubular structure and branching morphogenesis, like mammary glands (4). It has also just recently been recognized that biomechanical cues and mechanical interactions between cells also affect vascular development and organ specificity of vascular systems. This has, to date, largely been ascribed to shear stress exerted *via* blood flow (5). However, there are also numerous reports showing that endothelial cells respond to biophysical confinements (6), matrix stiffness (7, 8), and fibrous topography (9, 10). Furthermore, endothelial cells are able to deform collagen I gels (11) and thereby to communicate mechanically (12). This clearly indicates that biomechanical regulation of endothelial cells is much more versatile than only responding to shear stress. Unfortunately, so far, not much has been uncovered about the role of biomechanical processes in vascular development. Here, we aim to address this issue by experimentally probing the influence of biomechanics on cellular pattern formation. *In vivo* endothelial cells are mainly exposed to a vascular basement membrane (BM). Instead of collagen I this membrane consists of laminins and collagen IV (13), which form a non-fibrous matrix on a micron scale (14–16). One of the most widely used *in vitro* assays for angiogenesis, the “tube formation assay”, employs Matrigel, which is a reconstituted basement membrane (BM) prepared from a murine tumour (17). In this assay, single endothelial cells find each other within minutes and start to form a network. The mechanisms underlying this initial finding process and the subsequent pattern formation are still unclear. Here, we address this open issue and perform an in-depth analysis of the interactions between endothelial cells and Matrigel using functional assays, confocal microscopy and atomic force microscopy (AFM). We report that, while chemotactic gradients do not seem to play a role for the initial finding process between cells, the cells actively restructure the matrix. This leads to stiff fibrous bridges between nearby cells, which then become tracks for cell-cell contacts. In the light of recent studies showing matrix remodelling by diseases like fibrosis (18, 19), as well as therapeutical accessibility of the ECM (20), our data might help to understand the forces shaping vascular beds.

## Results

### The early finding phase of cells depends on cell density and has intrinsic length scales

After seeding the cells on Matrigel, cell-cell contacts are rapidly formed (Fig. 1A, Supplementary Movie S1). Subsequent to this initial “finding phase”, tubes are formed and re-organized. To investigate the impact of cell-cell distance on network formation, tube formation assays were performed with different cell densities. At low densities (up to 6 x 10^4^ cells/ml), the cells barely moved, and no network formation occurred (Fig. 1B). With higher cell numbers, tube formation was observed (Fig. 1B). To characterize cell behaviour during tube formation, we analysed a large body of individual trajectories obtained from cells at eight different seeding densities (from 0.5 to 20 x 10^4^ cells/ml). Since cells only move significantly when tube formation occurs, we concluded that cell motility must be a result of the macroscopic self-organization of the cells into a network, and not the other way around. Therefore, we directly related the behaviour of single cells to the behaviour of the collective and found the typical time scale of tube formation to be 93 min in the normalized velocity autocorrelation function (Fig. 1C, lower left panel)]. Furthermore, we observe network formation only above a critical cell density, which suggests that cells cannot detect each other if they are far apart. To test whether there is an intrinsic length for intercellular signalling, we first measured the mutual velocity alignment of distant cells. Here, we found that cells weakly align their motion across the whole field of view, and especially so within a typical radius of 106 µm (Fig. 1C, lower right panel). Finally, we confirmed that there is a directed component of cell motion, so that cells sense the positions of surrounding individuals. We found that cells are attracted to distant cells, while being sterically repelled from nearby cells. The distance over which cells optimally sense other cells coincides well with the distance over which cells align their direction of motion (Fig. 1C, upper right panel). Taken together, these findings suggest that cells can sense each other, to align and cluster, over a typical distance of at least 106 µm. To communicate over such long distances, cells would have to employ either chemical or mechanical signalling. Therefore, we next narrowed down which type of cellular signal is necessary for tube formation.

**Figure 1:**
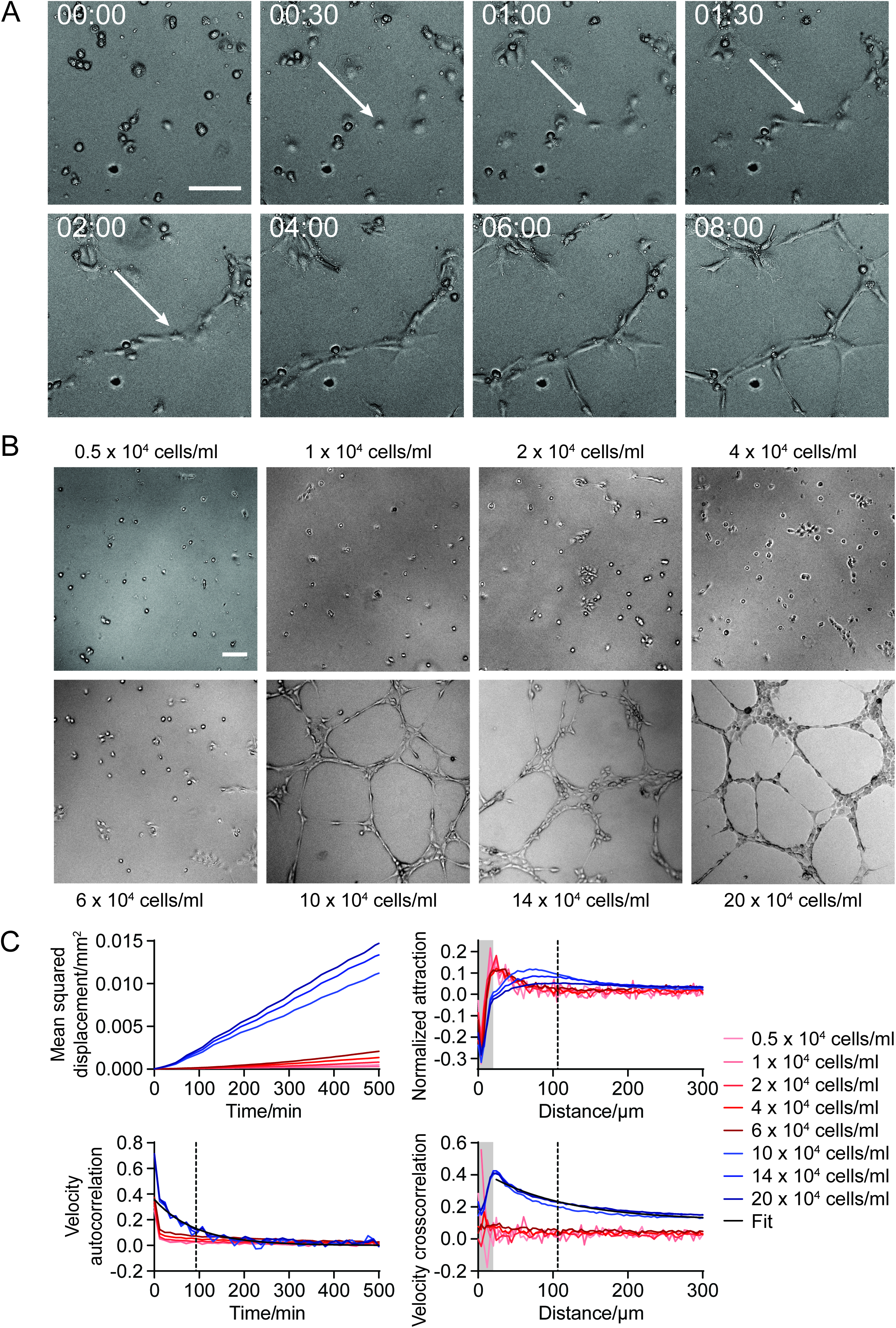
The initial finding phase depends on cell-cell distance. **(A)** HUVECs rapidly form cell-cell contacts when seeded on Matrigel. White arrows indicate the initial formation of the first cell-cell contacts in a time range from 1 to 2 h. Scale bar: 100 µm. **(B)** Tube formation after seeding different cell densities from 0.5 x 10^4^ cells/ml to 20 x 10^4^ cells/ml. At least 10 x 10^4^ cells/ml were necessary to induce tube formation, while at lower cell numbers, HUVECs stayed separate or formed small group of cells. Images were taken after 6 h. Scale bar: 100 µm. **(C)** The cell densities inducing tube formation (blue curves) showed a higher mean squared displacement compared to the cell densities without tube formation (red curves). The normalized attraction and the velocity cross-correlation showed that cells can sense each other, to align and cluster, over a typical distance of at least 106 µm (vertical dotted line in the attraction graph; the typical interaction distance is extracted from an exponential fit with a constant offset, which is indicated in black). The typical time scale for the tube formation with 93 min is obtained by fitting the velocity autocorrelation with an exponential (indicated in black). A typical cell size is indicated by the grey region.

### The initial phase of tube formation is not based on soluble or matrix bound chemotactic gradients

To investigate what causes the initial finding of cells and early pattern formation, we first examined the potential role of chemotactic gradients. To override endogenous gradients, we added 20 nM VEGF to the medium. After 3 h and 6 h we detected no significant difference in the number of tubes and nodes (Fig. 2A). Even when cells were totally deprived of growth factors by seeding them in phosphate buffered saline (PBS), initial tube formation took place (Fig. 2A). To specifically avoid the formation of soluble gradients in the medium, we also performed microfluidic experiments, where the tube formation setting was constantly superfused. However, tube formation still occurred (Fig. 2B and Supplementary Movie S2) – even though the tubes were distorted by shear stress.

**Figure 2:**
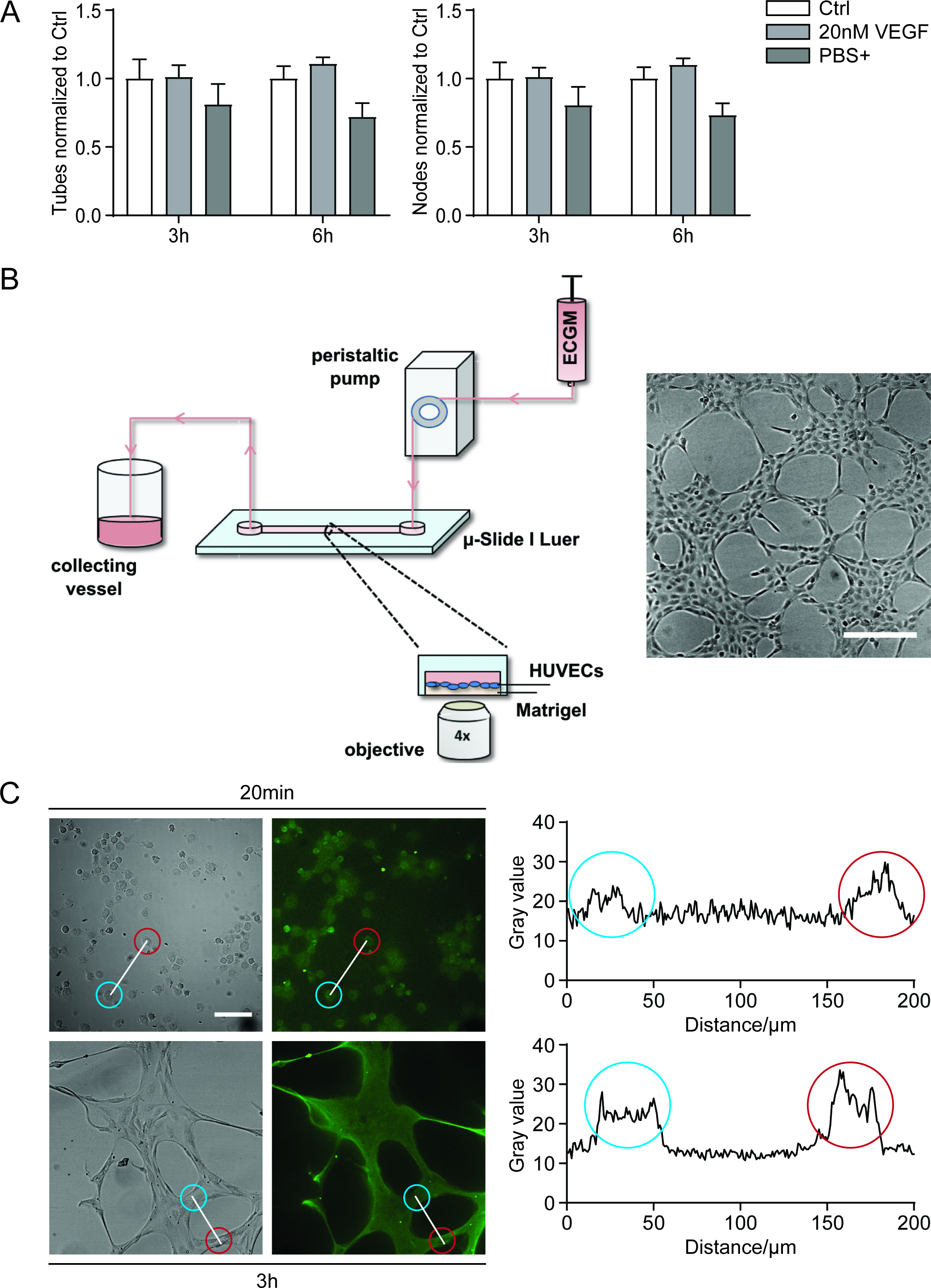
Tube formation does not depend on chemotactic gradients. **(A)** Quantification of the number of tubes and nodes after 3 h and 6 h. Neither a saturation of potential gradients by externally adding 20 nM VEGF, nor complete deprivation of growth factors (PBS+ instead of growth medium) significantly changed tube formation (One-way ANOVA, Dunnett’s test, compared to control). **(B)** To disturb potential local gradients of chemoattractants, the tube formation setting was continuously superfused in a microfluidic device (left panel). The absence of soluble gradients did not impair HUVEC tube formation (right panel, see also Supplementary Video 1). Scale bar: 100 µm. **(C)** To detect potential bound gradients of VEGF, an immunostaining for VEGF was performed (left panel). VEGF levels are highest around the cells, and no continuous long-ranged gradients between cells can be detected. Right panel: Fluorescence intensity profiles along the indicated white lines in the left panel. Scale bar: 100 µm. The read and blue circles around cells in the images correspond to the respective peaks in the intensity diagram.

Next, we investigated the potential existence of matrix bound gradients. As shown by immunostainings and intensity profiles (Fig. 2C), there was no matrix bound gradient of VEGF (green) between single cells, or tubular structures respectively. At the starting point right after cell adhesion (20 min), the fluorescence intensity of VEGF was evenly distributed with a slightly elevated intensity at the areas covered the cells, but without a long-ranged gradient. The intensity plot after 3 h showed an equal progression: The intensity of VEGF bound to the matrix was at a basal level, and there was an increased intensity at the areas covered by cells or tubes, respectively. However, there were still no apparent long-ranged VEGF gradients. In an additional set of experiments, we inhibited matrix metalloproteases (MMPs) with 10 µM batimastat to block the potential release of matrix bound growth factors. Batimastat treatment did not reduce tube formation (Supplementary Fig. S1). Therefore, we conclude that protein gradients are not the leading cause of tube formation.

### Tube formation depends on cell contractility, matrix deformation/remodelling and matrix stiffness

To test the hypothesis of mechanical signalling as driver of tube formation, we treated cells with the myosin II ATPase inhibitor blebbistatin. This caused a loss of contractility, and initial network formation was dramatically reduced (Fig. 3A). Next, we performed traction force microscopy experiments to investigate matrix deformation during tube formation (Fig. 3B). The analysis of the bead displacement over the first 3 h after seeding the cells showed significant matrix deformation. Matrigel was detectably compressed in a distance up to only 30 µm around the cell, but the displacement of the beads could be detected at distances around at least 100 µm. This nicely corresponds to the normalized attraction between cells detected by cell tracking (Fig. 1C). With blebbistatin treatment the deformation was weaker and slower, but the length scale of substrate compression remained roughly the same.

**Figure 3:**
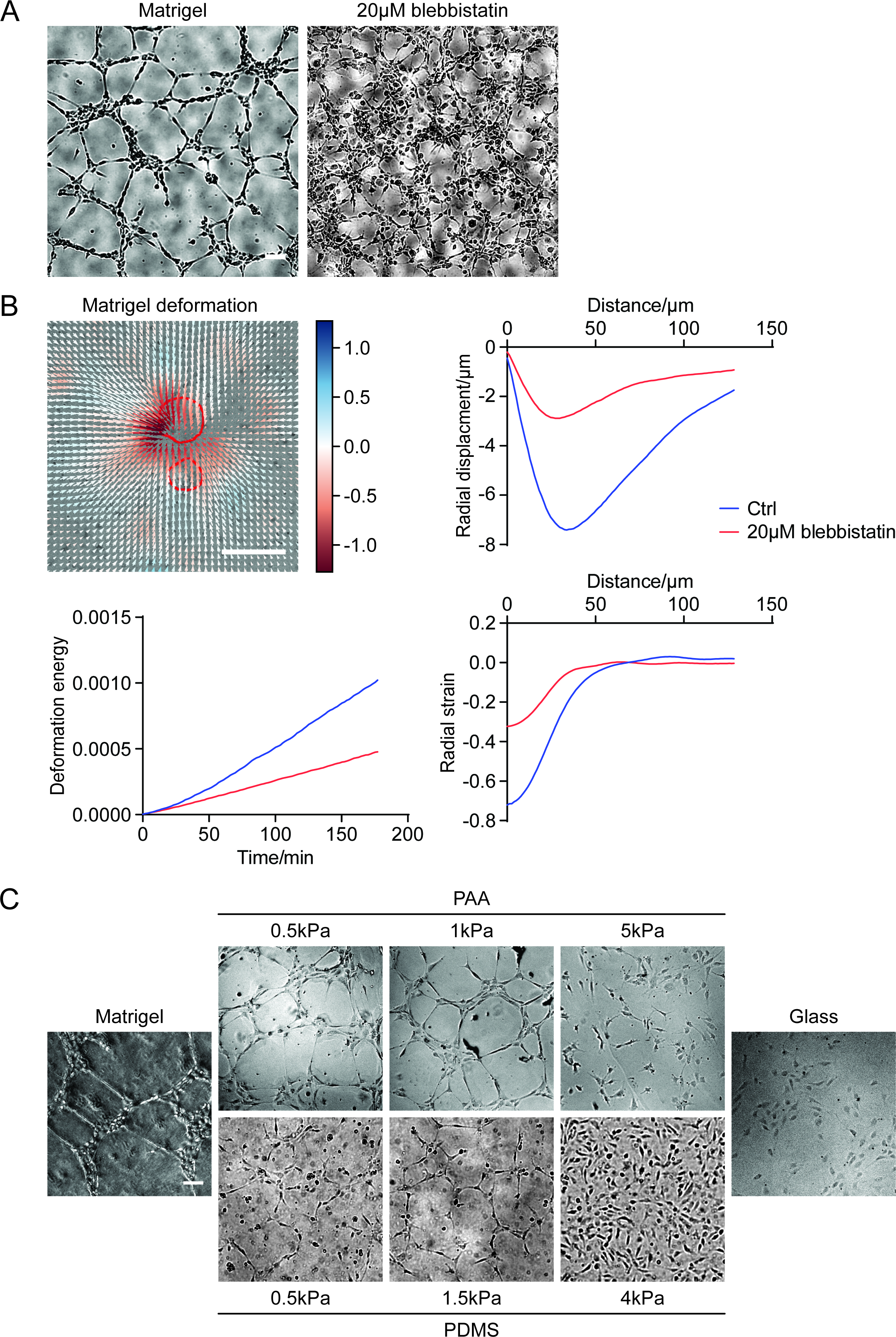
Tube formation depends on active matrix deformation and substrate stiffness. **(A)** Inhibition of cell contractility with 20 µM blebbistatin inhibited the formation of tubular structures. Scale bar: 100 µm **(B)** Upper left: Representative image of bead displacement by cells (red circles) on Matrigel as a dimensionless heatmap. Upper right: The analysis of the bead displacement over the first 3 h after seeding the cells on Matrigel showed that cells deform the matrix with a maximum around 40 µm. The energy 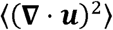 (lower left) and radial strain (lower right) **(C)** HUVECs treatment with 20 µM blebbistatin reduced radial displacement, bulk deformation (20 x 10^4^ cells/ml) were seeded on PAA or PDMS gels with different stiffness from 0.5 kPa to 5 kPa, coated with Matrigel, on Matrigel alone or on glass coated with Matrigel. At high (4 kPa and glass) stiffness values, the cells formed no network. On substrates with a stiffness between 0.5 and 1.5 kPa, HUVECs formed a tubular network like on the Matrigel control. Scale bar: 100 µm.

Since matrix deformation seems to play an important role for tube formation, we created polyacrylamide (PAA) and polydimethylsiloxane (PDMS) gels of defined stiffness and coated them with a layer of Matrigel of defined thickness (< 20 µm). This setup allows for sensing of the underlying polymer surface by the cells (Supplementary Figs. S2 and S3). At stiffness values between 0.5 kPa (similar to Matrigel) and 1.5 kPa HUVECs form a tubular network on both polymers, comparable to the Matrigel control (Fig. 3C). At a stiffness of 4 or 5 kPa, HUVECs just spread like on the glass control (Fig. 3C). When we stained and imaged the two main constituents of Matrigel, laminin and collagen IV, we observed a dramatic remodelling over time (Fig. 4): the cells rearranged laminin and collagen IV, forming bridges of matrix material between them. Inhibiting cell contractility via blebbistatin treatment strongly inhibited this behaviour (Fig. 4) so that no bridges were observed. We find that matrix remodelling depends on matrix stiffness: while cells remodelled the matrix at low stiffness values (0.5 and 1.5 kPa), a stiffer gel (4 kPa) stayed largely unstructured (Supplementary Fig. S3C). This suggests that the ability to deform and restructure the matrix is crucial for subsequent tube formation, and depends on cell contractility and matrix stiffness.

**Figure 4:**
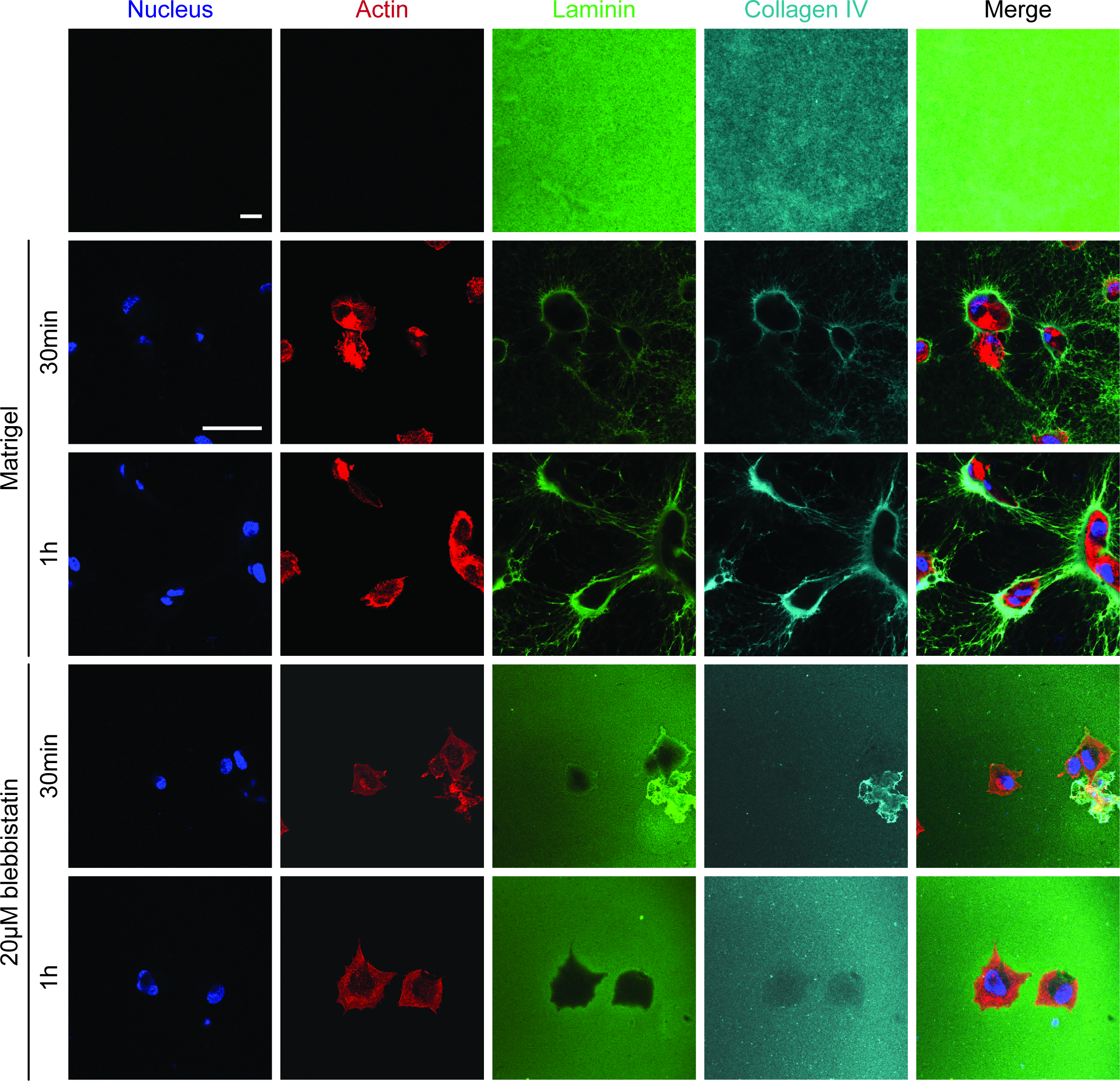
Endothelial cells actively remodel ECM during early tube formation. Immunostaining of laminin and collagen IV in Matrigel without (first row, scale bar 10 µm), and with cells (second and third row, scale bar 50 µm). Cells time dependently re-arrange the matrix proteins so that fibres between cells are formed. This process depends on cell contractility and is inhibited by 20 µM blebbistatin (fourth and fifth row).

### Laminin plays a key role for the tube formation

To investigate, whether substrate stiffness alone is the key parameter for initial tube formation, we coated PDMS gels with a collagen I gel to obtain the same stiffness as with Matrigel. In this case, the HUVECs spread without forming tubes (Supplementary Figs. S4A and B, and S5A). Most likely, this difference between Matrigel and a collagen gel of identical stiffness originates from the respective molecular and structural makeup of both gels. To investigate this point, we first focussed on laminin, the main constituent of Matrigel. A commercially available pure laminin gel turned out to be much softer that Matrigel (Supplementary Fig. S4A), but supported the initial tube formation process. At later time points, however, the tubular networks collapsed (Supplementary Fig. S4B). This suggests that the soft laminin gel could not withstand the contractile forces of cell collectives. When we created mixed gels with different ratios of laminin and collagen I, only the ratio of 6:1 (4.8 mg/ml:0.8 mg/ml, other ratios not shown) supported tube formation over a longer period of time (Supplementary Fig. S4B and C), although this gel was as soft as laminin alone. Thus, the presence of laminin seems to be of crucial importance. The presence of small amounts of fibrillary collagen I seems to lend mechanical stability to the gel, while a ratio above 6:1 disturbs the matrix reorganization process.

Next, we manipulated the structure of Matrigel with netrin-4, which binds to laminin-γ1 and is able to destroy laminin networks (21). We tested ratios between netrin-4 and laminin of 1:2, 1:1 and 1.5:1 (Fig. 5A). With a laminin excess no change in the laminin structure was observed. Both in pure Matrigel and for a netrin-4 to laminin ratio of 1:2, laminin showed a very homogeneous structure and cells formed a tubular network. For a netrin-4 to laminin ratio of 1:1, the homogenous structure of laminin was slightly interrupted, and the cells formed clusters instead of a network. With a netrin-4 excess, the structure of the laminin network completely changed and became rough (Fig. 5A). HUVECs seemed to sediment through the gel and spread on the plastic bottom of the well. Accordingly, not only the presence of laminin is of importance for tube formation, but also the crosslinking of laminin and the homogeneity of the substrate.

**Figure 5:**
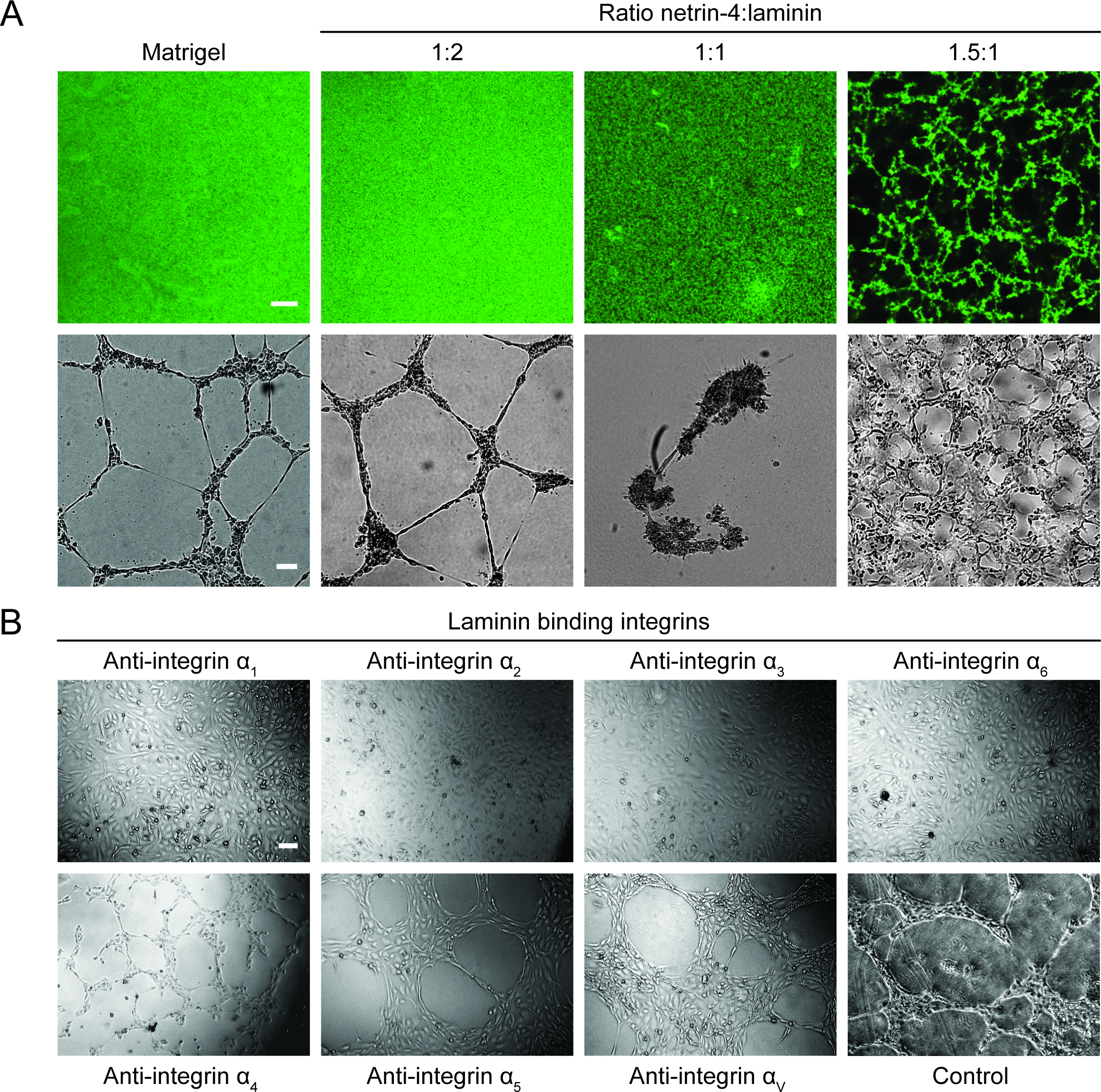
Laminin is a key factor for tube formation. **(A)** Upper row: immunostaining of laminin in Matrigel alone or after treatment with different rations of netrin-4 (Scale bar: 10 µm). Lower row: Tube formation on Matrigel and with different ratios of netrin-4 to laminin. Starting from a ratio of 1:1, netrin-4 changes the structure of the laminin network and influences tube formation. **(B)** Functionally blocking antibodies against integrins α1, α2, α3, α4, α5, α6 and αv were added (40 µg/ml). Blocking integrins, which preferentially bind to laminin (α1, α2, α3 and α6) inhibited tube formation, while blocking of mainly collagen and fibronectin binding integrins (α4, α5 and αv) did not affect endothelial tube formation. Scale bar: 100 µm.

As a different approach to test the relevance of laminin for tube formation, we used integrin-blocking antibodies (Fig. 5B). While inhibition of integrins favouring binding to fibronectin and other ECM proteins (integrins α_4_, α_5_ and α_v_) did not influence endothelial tube formation, blocking of integrins that mainly bind to laminin (integrins α_1_, α_2_, α_3_ and α_6_), potently inhibited the formation of tubular structures. This again underscores that laminin is the crucial matrix component for the formation of tubes on Matrigel.

### Strain stiffening guides the cell-cell sensing process

To get an impression of the mechanical landscape of the different matrices in the absence and presence of cells, we prepared stiffness maps by using atomic force microscopy (AFM). In collagen I gels we detected an inhomogeneity of the stiffness map, with alternating stiff regions at the collagen fibre locations and soft regions in between (Fig. 6A). In contrast, we found that the stiffness of Matrigel was homogeneous across the whole gel. By using fluorescence reflection microscopy, we could clearly detect fibres in collagen gels (Fig. 6A, right panel), while Matrigel showed no fibrous structure (not shown). On the collagen gels, the cells aligned along the stiff collagen fibres (Fig. 6B). The cell tracking plots shows that on collagen, single cells persistently migrated over long distances. In contrast, on Matrigel, single cells without interactions with other cells moved only over very short distances (Fig. 6B). Using AFM, we measured the stiffness of the areas between two cells (below the critical cell-cell distance for tube formation). As a control, an area not affected by cells was also measured. For all cell pairs, we could detect a significant increase in the stiffness of the space between cells, just before cells protrude into this area (Fig. 6C). It seems like the cells use the self-created stiffer regions as a guidance cue. Supplementary Fig. S6 shows further examples for the strain stiffening effect and its use for connecting cells.

**Figure 6:**
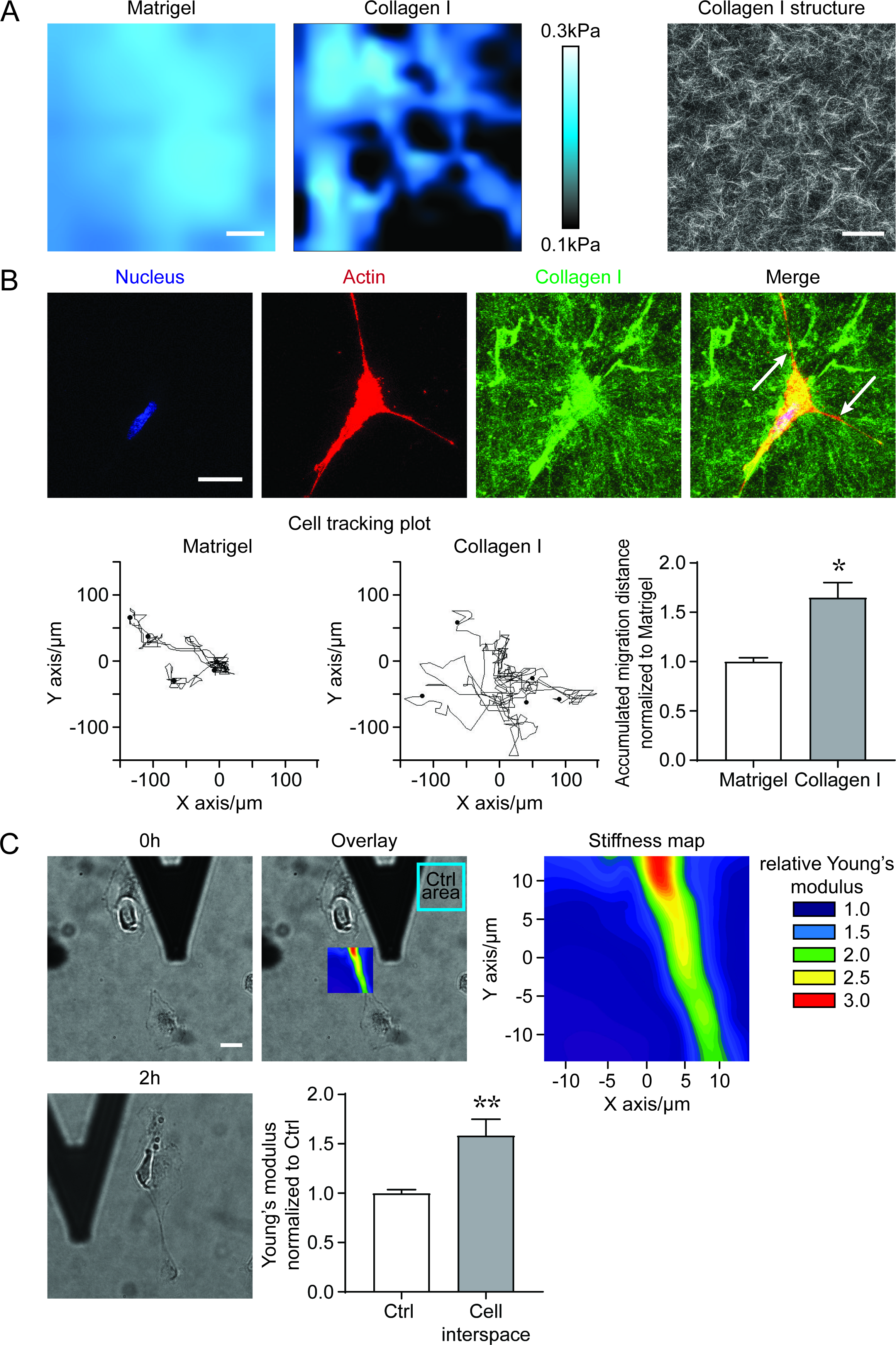
Cells change the local stiffness topography of the substrate. **(A)** Pseudo coloured AFM stiffness map of Matrigel and rat tail collagen I gel (Scale bar 5 µm): while the stiffness is evenly distributed in Matrigel, the collegen gel contains stiff fibres and soft interspaces. Right panel: the total reflection microscopy image of the collagen I gel shows the collagen fibre network (scale bar: 100 µm). Matrigel shows no fibrillary structure (data not shown). **(B)** Immunostaining of a collagen I gel in the presence of an endothelial cell. White arrows mark collagen fibres to which the cell is aligned (scale bar: 25 µm). Exemplary cell trajectories on Matrigel and collagen I over 20 h. Single cells on collagen migrated over a longer distance in all directions. The accumulated distance for 45 cells for each gel is normalized to the Matrigel and showed a significant increase on collagen I. Two-tailed unpaired Student’s t test with Welch’s correction, *P < 0.033. **(C)** The stiffness of the area between two single cells after adhesion on Matrigel was measured with an AFM and compared to the stiffness of a cell-free part of the gel. The pseudo-coloured stiffness map shows the relative Young’s modulus normalized to the control area. From the overlay of the microscope image and the stiffness map it can be seen that the cells formed a stiff bridge between them, before cell protrusions cover this area. Over all cell pairs (n=21) we detected a significant increase in the Young’s modulus of the cell interspace. Two-tailed unpaired Student’s t test with Welch’s correction, *P < 0.002 Scale bar: 20 µm. See also Supplementary Fig. S6.

## Discussion

Since the development of Matrigel, an extracellular matrix preparation from murine tumors, by the group of Hynda Kleinman nearly 30 years ago (22), the endothelial tube formation assay has become one of the most widely used *in vitro* assay systems for investigating angiogenesis (23). It is highly intriguing to note that the driving force behind the self-organization of the tube-forming cells is still unclear. The common assumption is that chemotactic gradients are the main sculptors of vascular development. VEGF is undoubtedly a central player in the processes of angiogenesis and vasculogenesis (1), since it guarantees endothelial cell survival, proliferation and chemotaxis. Furthermore, it coordinates with other signalling pathways (e.g. Notch), which are important for the differentiation of endothelial cells into tip and stalk cells (24). However, in our experimental model we could exclude that soluble gradients of VEGF play a role for the initial sensing phase of tube formation. Furthermore, matrix bound gradients of VEGF, which have been demonstrated in collagen gels (25), were not detectable in Matrigel during tube formation. This does of course by no means indicate that VEGF is not important at later time points (26) or for navigating new vessels into hypoxic tissues, but points to additional, alternative pathways of endothelial cell communication, like, e.g. mechano-signalling. We and others see that tube formation relates to cell density (26) and stiffness of the matrix (8). We hypothesize that endothelial cells generate deformation fields in the matrix, which allow for communication over longer cell-cell distances. A further hint towards a mechanical component during pattern formation is the drastic reduction of tube formation after inhibition of cell contractility *via* blebbistatin treatment.

In fibrous gels (mostly collagen) mechanical sensing between cells over long distances (approx. 100 µm) has been repeatedly described and has been ascribed to two phenomena: fibre alignment and strain stiffening in response to cellular traction forces (27, 28). Furthermore, it has been shown that cell-induced matrix stiffening is more profound for cell ensembles than for single cells (10, 29). Similar observations have also been made in fibrin gels (30, 31). On linearly elastic materials, like polyacrylamide gels, the range of cellular force sensing was much smaller (32). This leaves two major open questions concerning our experimental data: 1) How do cells communicate mechanically in a non-fibrous gel like Matrigel? and 2) Why are the cells not able to form a network on fibrous (collagen) gels?

Matrigel mainly consists of laminins, which are of primary importance for tube formation (17, 22). In fact, functionally blocking antibodies against integrins that mainly bind to laminin inhibits tube formation, while blocking other integrins has no significant effect. While laminin gels are *per se* amorphous, it has been shown that cell-sized fibrous structures can be induced by mechanical forces, like the external application of fluid flow (33) or exerting a force via paramagnetic particles in a magnetic field (34). Here, we observe that Matrigel is rearranged into fibres by the exposure to cells. Since this phenomenon occurs rapidly and at intercellular regions, which have not been covered by cellular protrusions, we can exclude that this is due to secretion of fibrillary proteins. Furthermore, mixing netrin-4, which has been shown to disrupt laminin networks (21), into Matrigel preparations inhibited tube formation. This underscores that rearranging laminins into fibrous structures via traction stresses is a fundamentally important feature of tissue organization. Similar phenomena might occur *in vivo*, since extracellular spaces where vasculogenesis takes place during development (35, 36), or angiogenesis during tissue regeneration (37) are generally rich in laminin.

Rearrangement of extracellular matrix can influence cell behaviour in multiple ways. Recently, it has been shown that cells can sense the density of fibre networks, which was termed “topotaxis” (41). At high pre-existing fibre densities (like in a conventional collagen I gel) cells stay unorganized. At lower densities of pre-formed fibres (but still the right stiffness of the gel), endothelial cells have recently been shown to align to these fibres (9). At a low cell density, we make a similar observation on collagen: cells align to preformed fibres. However, unlike on Matrigel, they do not organize themselves into tubes. Furthermore, Matrigel – similar to collagen or fibrin gels – shows stress stiffening after cell contraction (38). As a consequence, cells could migrate along the ensuing stiffness gradient, a phenomenon called “durotaxis” (39, 40). Then why do endothelial cells form tubes on Matrigel by inducing a fibrous structure in their environment, but not on collagen I gels of similar stiffness, which are fibrous from the start? A possible hint lies in the different behaviour of single cells and cell collectives: though the single cells in our case were not able to communicate on collagen I gels, endothelial spheroids have been shown to sense each other over large distances in collagen gels (10). On Matrigel, even single cells are able to restructure the previously homogenous matrix into a fibril network, where fibres are aligned between cells. The resulting rigid structures might be the basis for durotaxis, topotaxis, or might be places of high adhesive ligand density (for a schematic overview see Fig. 7). On collagen, single cells are not able to do induce a significant restructuring as collagen is already inhomogeneous from their perspective. However, on the length scale of an organoid, collagen might seem almost like a homogeneous environment. This might explain why organoids are able to communicate via restructuring their extracellular matrix, where single cells are not able to do so.

**Figure 7:**
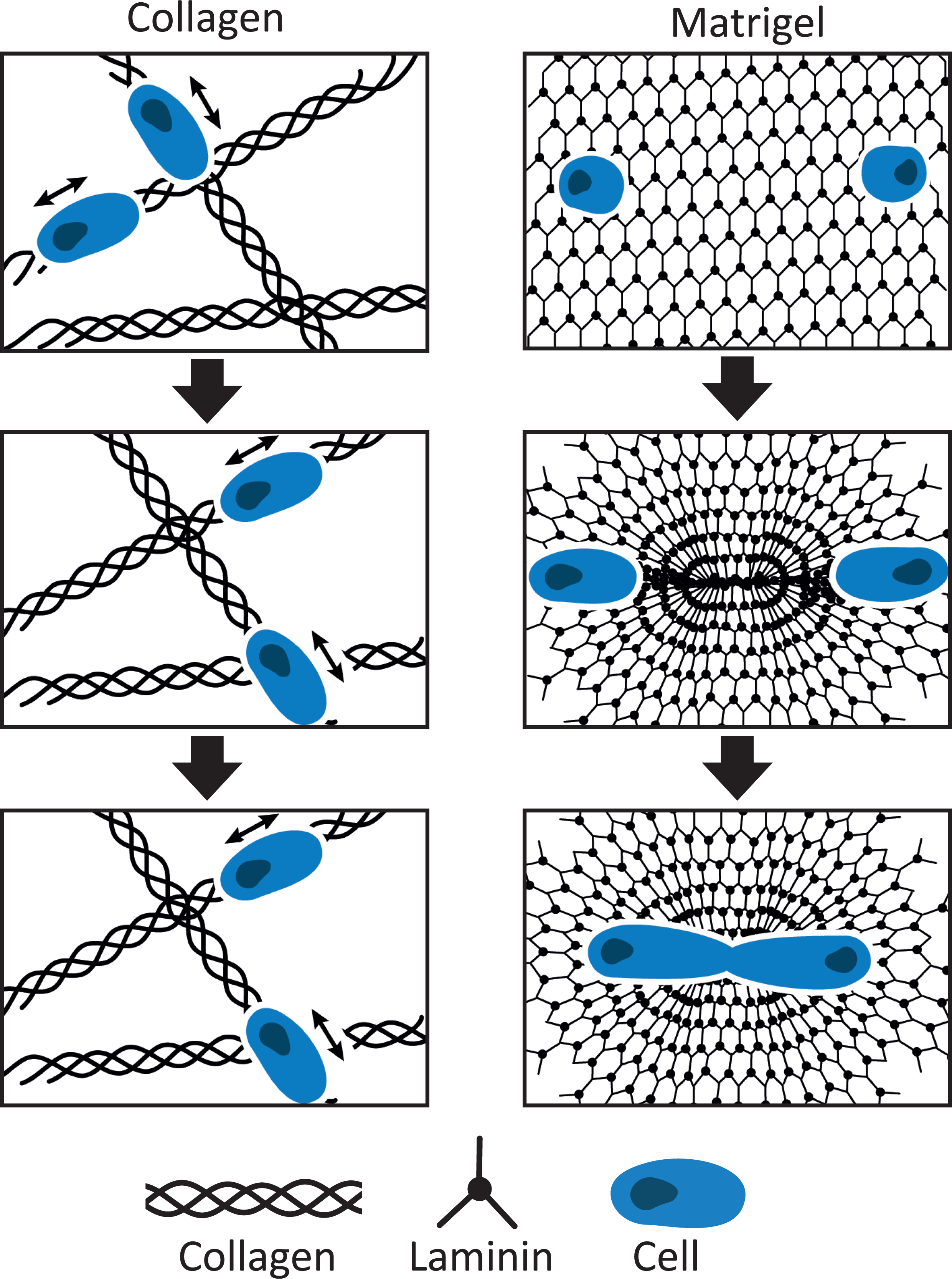
Graphical Summary. While endothelial cells align to collagen fibres and move along them (left panels), they remodel the laminin based Matrigel (right panels): by contractile forces, strain stiffening occurs and the stiffened intercellular bridges are used as tracks for pattern formation.

Accordingly, our concepts of factors shaping a vascular network need to be extended: Though chemotactic gradients are doubtlessly important, they are definitely not the only key. Tissue and matrix stiffness, biochemical constitution (e.g. the presence of laminin or collagen I), as well as topographic cues from the matrix (is the matrix homogenous or heterogeneous?) play important roles. The latter ones might rely on physical differences: in homogeneous gels, deformation fields are transmitted across wide area, while fibrous gels transmit forces in a rope-like fashion. From a systemic perspective, the high variability of extracellular matrices throughout the tissues of the body, which is even remodelled by diseases like fibrosis (18, 19), might be the basis of highly flexible and adapted vascular networks. The apparent importance of the mechanical properties of the ECM, as well as its constituents, for morphogenesis and pattern formation gives access to an exciting new field, which might even give rise to new therapies of metabolic diseases/cancer. Similar concepts might also apply for morphogenic processes in other contexts.

## Material and Methods

### Cell culture

HUVECs were purchased from Promocell (Heidelberg, Germany) and maintained in EC growth medium ([PB-MH-100-2199] Pelobiotech, Planegg, Germany) containing, 10% fetal calf serum (FCS), 10.000 U/ml penicillin/streptomycin and 250 µg/ml amphotericin B under constant humidity at 37 °C and 5 % CO_2_. Experiments were performed using cells at passage 6.

### Tube formation assay on Matrigel

HUVEC tube formation assays were performed in µ-Slide angiogenesis or µ-Slide 8 well (ibidi, Martinsried, Germany). Matrigel (Corning, Amsterdam, Netherlands) was thawed on ice over night the day before use. For homogeneity the gel was mixed thoroughly after thawing and kept on ice. Where indicated, Netrin-4 (a kind gift from R. Reuten) was mixed with Matrigel at ratios of 1:2, 1:1 and 1.5:1 netrin-4 to laminin. 10 µl and 30 µl of Matrigel were used to fill the inner well of the angiogenesis slides or 8 well slides, respectively. For gelation, the gel was kept at 37 °C, 5 % CO_2_ for at least 30 min. After reaching confluency, HUVECs were trypsinized and diluted to the desired density with EC growth medium. Compounds (blebbistatin; batimastat, Sigma Aldrich, St. Louis, USA; α integrin blocking kit, Millipore, Darmstadt, Germany) for treatment were diluted with the cell suspension to the indicated final concentration. 50 µl of the cell suspension was applied to the upper well of the angiogenesis slide, and for the 8 well slide 250 µl of the cell suspensions were added. The slides were incubated at 37 °C and 5 % CO_2_, and Live imaging was performed on an inverted microscope Eclipse Ti (Nikon, Düsseldorf, Germany), with a 4x/10x phase contrast objective and a CCD camera ([DS-Qi1Mc] Nikon, Düsseldorf, Germany). The slide was inserted into a 37 °C heating and incubation system, which was flushed with actively mixed 5 % CO_2_ at a rate of 10 l/h and the humidity was kept at 80 % to prevent dehydration. Images were taken after a desired period of time on a Leica DMi1 microscope (Leica Microsystems CMS GmbH, Wetzlar, Germany). All images for the tube formation were analyzed using the ImageJ software (National Institutes of Health, Bethesda, USA) tool “Angiogenesis Analyzer” (Gilles Carpentier, Faculte des Sciences et Technologie Universite Paris Est Creteil Val-de-Marne, France) according to the number of tubes and nodes.

### Microfluidics

For culture under flow conditions the µ-Slides I Luer (ibidi, Martinsried, Germany) were coated with Matrigel (100 µl per Slide) and kept at 37 °C, 5 % CO_2_ for at least 30 min. HUVECs (5 x 10^5^ per cm^2^) were seeded into the channels with a micropipette, and the cells were allowed to attach to the gel for 20 min under static conditions at 37 °C, 5 % CO_2_. Afterwards the channel slide was inserted into an incubation system and placed on an inverted microscope. The humidity was kept at 80 % to prevent dehydration. Fresh EC growth medium was then perfused through the channel at a flow rate of 0.1 ml/min for 6 h.

### Tube formation assay with different cell densities

For cell tracking in the live imaging HUVECs were stained with Hoechst 33342 (Sigma Aldrich, St. Louis, USA). The confluent cells were washed with PBS^+^ (with Ca^2+^ and Mg^2+^) and then stained with 5 µg/ml Hoechst for 20 min. The tube formation assay was performed according to the protocol described above.

The positions of the stained cell nuclei were detected from the fluorescent channels of the tube formation videos, and then stitched together into sets of time-lapse cell trajectories. Here, we proceeded as follows. First, to detect the cell cores, we (i) improved contrast via a contrast-limited adaptive histogram equalization using *OpenCV* (https://github.com/opencv/opencv/wiki/CiteOpenCV), (ii) normalized all image data, and (iii) applied a multi-pass Laplace-of-Gaussian filter using *scikit-image* (https://scikit-image.org). Then, we linked the individual cell core coordinates into cell trajectories with *Trackpy (*https://zenodo.org/record/1226458). An alternative analysis with the software *ImageJ* and its plugin *TrackMate* (42) yielded similar results (not shown). Further analysis of the obtained cell trajectories was performed, as described below.

To measure whether cell motion during tube formation could be described by a persistent random walk with a typical timescale, a measure for the correlation between the direction of cell motion at a reference time *t*, and a time, +Δ*t*, the normalized velocity autocorrelation function, was introduced:

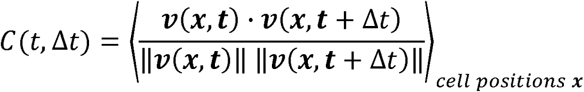

For a time-independent process, one can average over all available reference times *t*,. Tube formation, however, is a time-dependent process which might be subject to aging, for example if cell velocities change over time. Therefore, here we chose to average over reference times that lie within a window, 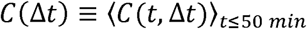. This time window is shorter than the timescale of tube formation (which we show to be 93 *min*), and much shorter than the overall duration of the experiments (20h). We found that the normalized velocity autocorrelation function decays rapidly from *C*(0) = 1 to *A* = C(10 *min*) ≈ 0.42 after the first frame. For all subsequent frames, the typical timescale of tube formation, *τ*, was obtained from an exponential fit: *C*(Δ*t*) ≈ *A exp*(-Δ*t/τ).*

A possible correlation between the migration of distant cells, as a function of separation distance between the considered cells, was determined as follows:

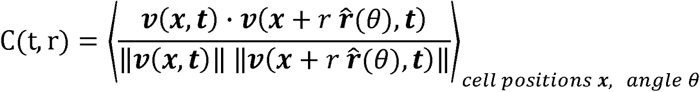

Here, we measured the cosine of the angle between the velocity vector of a cell at position *x*, and a cell at position *x+r θ,* where r(*θ*) is the radial unit vector pointing in the direction of the angle *θ* (in cylindrical coordinates). Again, we averaged over reference times that lie within a time window, 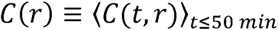. Then, the typical length scale of cell-cell velocity alignment *r*_1_, was obtained from a bi-exponential fit: *C*(*r*) ≈ *A exp*(-*r*/*r*_1_)+*B exp*(-*r*/*r*_2_), where *r_1_* <<*r_2_*

Finally, a correlation between the migration of a cell and the position of a distant cell that lies in the direction r(*θ̂*), as a function of their separation distance r, was determined in a similar way:

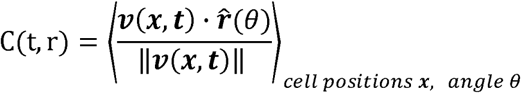

Note that this measure does not vanish if the cells are distributed inhomogeneously.

### Traction force microscopy

For the traction force microscopy HUVECs were prepared according to the protocol for the tube formation assay on Matrigel with µ-Slides 8 well. Yellow-green microbeads (Life Technologies, Carlsbad, USA) of 1 µm diameter, 2 % solids were mixed with Matrigel (1:200). To determine the forces of single cells, cells were seeded at a concentration of 10,000 cells/ml. Time-lapse video microscopy was performed with a time interval of 1 min between images over 3 h.

Substrate displacement fields, *u*(*x,t*), were obtained from the fluorescent channels of the videos by computing the corresponding optical flow fields using the “Two-Frame Motion Estimation Based on Polynomial Expansion” by Gunnar Farnebäck implemented in *OpenCV*. Then, the compression fields, 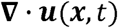, were determined and visualized in the phase contrast channels of the videos. We found that each (round) cell generated a radially symmetric compression field. Therefore, we determined the average radial component of the compression field, 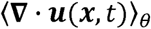, and the radial component of the displacement field, 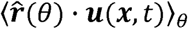, in the reference frame of the individual cells.

### Tube formation assay on other gels

HUVEC tube formation on rat tail collagen I gels (Corning, Amsterdam, Netherlands) was prepared according to the protocol for tube formation assays on Matrigel. The collagen I gel (2 mg/ml) consisted of 10 % 10x PBS, 1 N NaOH (0.023 times of the final volume of collagen I) and dH_2_O to adjust the collagen concentration. 10x PBS, 1 N NaOH and dH_2_O were mixed and kept on ice. Subsequently, the calculated volume of collagen I was added. For creating the laminin-collagen I mixture gel in a ratio 6:1 (4.8 mg/ml:0.8 mg/ml), a 4 mg/ml collagen I gel (produced according the protocol) and a 6 mg/ml laminin gel (Trevigen, Gaithersburg, USA) were mixed on ice. For gelation, the gels were kept at 37 °C, 5 % CO_2_ for at least 1 h. Organ specific ECM gels were from Xylyx (New York, USA) and used according to the manufactureŕs protocols.

### Polyacrylamide gels coated with Matrigel

Elastic polyacrylamide gels with a Young’s modulus from 0.5 to 5 kPa were prepared as reported elsewhere (43). For cell adhesion, Matrigel was covalently attached to the polyacrylamide gels and to amino-silane coated glass with the bifunctional cross-linker Sulfo-SANPAH (Thermo Fisher Scientific, Waltham, USA), respectively. HUVEC tube formation assay was performed as described before.

### Gel printing on polydimethylsiloxane (PDMS)

In order to determine the maximum thickness of Matrigel, which still allows cells to feel the stiffness of an underlying PDMS gel (Sylgard 184 silicone elastomer kit, Dow Corning, Midland, USA), different volumes were tested in the angiogenesis slide. PDMS and the curing agent were mixed and dried in a desiccator for 15-20 min in order to remove air bubbles. Afterwards, the gels were baked for 20 h in a compartment drier at 60 °C. The inner well of the angiogenesis slide was filled with a very stiff PDMS gel (10 % curing agent), which, as such does not support tube formation. PDMS was hydrophilized with oxygen plasma for 3 min at 0.3 mbar in a plasma cleaner (Diener Plasma-Surface-Technology, Ebhausen, Germany). Matrigel was mixed with 5(6)-FAM, SE (5-(and-6)-Carboxyfluorescein, Succinimidyl Ester) (Thermo Fisher Scientific, Waltham, USA) with a final concentration of 0.1 mg/ml and volumes from 10 µl to 1 µl were added on the PDMS. The thickness of the Matrigel layer was measured on a Leica TCS SP8 confocal microscope (Leica, Wetzlar, Germany) with HC PL FLUOTAR CS2 10x/0.3 NA DRY using LAS X Core Software (Leica, Wetzlar, Germany). Even at a layer thickness of 25 μm, the cells formed a network and did not feel the stiffness of the underlying PDMS (Supplementary Fig. S2). To obtain an even thinner layer, PDMS was filled into a 2 well slide (ibidi, Martinsried, Germany), then hydrophilized, and 5 µl of Matrigel was added. A PDMS stamp was pushed on the Matrigel to equally distribute it (model in Supplementary Fig. S3A). This procedure allows a layer thickness under 20 µm, which is thin enough for the cells to sense the stiffness of the underlying PDMS, because they did not form a network any longer (Supplementary Fig. S2). PDMS and the curing agent were mixed in different ratios in order to obtain various gel stiffnesses. PDMS gels with a Young’s modulus of 0.5 (1.3 %), 1.5 (2 %) and 4 kPa (2.22 %) were prepared (Supplementary Fig. S3B). The stiffnesses of the PDMS gels were measured after the hydrophilization with a Modular Compact Rheometer MCR 100 (Physica, Stuttgart, Germany) in the amplitude sweep mode with a constant frequency of 1 Hz at 37 °C between PP25 measuring plate. The deformation was varied from 0.01 % to 10 % in a ramp mode. The measurements were averaged after applying the Grubbs outlier test. For the gel printing, Matrigel and collagen I gel (2 mg/ml) were used.

### Immunostaining and confocal microscopy

Cells were washed with PBS for 10 min and fixed after desired time points with 4 % para-formaldehyde for 10 min, followed by three washing steps with PBS for 10 min, and blocking with 1 % bovine serum albumin (BSA) in PBS at least 2 h or overnight at 4 °C. After the blocking step the slide was washed with PBS for 10 min. The primary antibodies (anti-VEGF (rabbit) 1:100 dilution [ab46154] Abcam, Cambridge, UK; anti-collagen I (rabbit) 1:200 dilution [34710] Abcam, Cambridge, UK; anti-collagen IV (rabbit) 1:100 dilution [AB756P] Sigma Aldrich, St. Louis, USA; anti-laminin (rat) 1:200 dilution [MA1-06100] Thermo Fisher Scientific, Waltham, USA) were incubated overnight at 4 °C. Prior to incubation with the secondary antibody (Alexa Fluor 488 goat anti-rabbit 1:200 dilution [A-11008] Life Technologies, Carlsbad, USA; Alexa Fluor 488 goat anti-rat 1:200 dilution [A-11006] Life Technologies, Carlsbad, USA; Alexa Fluor 647 chicken anti-rabbit 1:200 dilution [A-21443] Life Technologies, Carlsbad, USA), the slide was washed six times with PBS for 10 min and then incubated overnight at 4 °C. Again, cells were washed three times with PBS for 20 min and stained with Hoechst 33342 in a final concentration of 5 µg/ml and rhodamine-phalloidin ([R415], Sigma Aldrich, St. Louis, USA) diluted with 1 % BSA in PBS for 1 h. Afterwards the preparation was washed three times with PBS for 10 min and mounted with FluorSave™ Reagent (Merck, Darmstadt, Germany). Confocal images were collected using the Leica TCS SP8 confocal microscope with HC PL APO CS2 20x/0.75 NA IMM, or 63x/1.4 NA OIL objectives using LAS X Core Software.

### Atomic force microscopy (AFM)

400 µl of the respective gel was distributed in a 40 mm petri dish. The petri dish was filled with EC growth medium and kept at 37 °C for the measurement. The AFM measurements were performed on a NanoWizard® 4 (JPK Instruments, Berlin, Germany) with an integrated Axiovert 200 inverted microscope (Zeiss, Jena, Germany) using SPM software (JPK Instruments, Berlin, Germany) in the contact mode. For measuring the gel stiffness homogeneity a MLCT-C cantilever (silicon nitride, resonance frequency 7 kHz, spring constant 0.01 N/m, Bruker, Berlin, Germany) was used and calibrated with the contact free method. The following values have been set: setpoint 1 nN, z-length 8 µm, speed 2 µm/s and pixel size 16×16 on a 30×30 µm grid. For measuring the strain stiffening, a tipless MLCT-D cantilever (silicon nitride, resonance frequency 15 kHz, spring constant 0.03 N/m, Bruker, Berlin, Germany) glued with a hollow glass bead ([PBGH-18] Kisker Biotech, Steinfurt, Germany) was used and calibrated with the contact free method. Immediately before the measurement, the diameter of the bead was measured. For the setting the following parameters were used: setpoint 2 nN, z-length 10 µm, speed 10 µm/s. The measuring area was adjusting to the distance of the cells. The pixel size was 10×10 and every pixel was measured three times directly successively. For the control area a 5×5 µm grid far away from the cells was measured after every cell measurement.

Data processing was performed using the corresponding software version 6.0.50 (JPK Instruments, Berlin, Germany). The Hertzian contact model (Young’s modulus) was used to calculate the stiffness. In the software the spring constant and sensitivity of the cantilever were loaded and then the baseline and contact point were determined. The tip shape of the MLCT-C was modelled as quadratic pyramid and the half-front angle of the cantilever set to 15°. For the MLCT-D with the bead a sphere model was used and the radius of the bead was set. The Poisson ratio was set to 0,5.

### Statistical analysis

Results of at least three independent experiments (biological replicates, each performed in two or three technical replicates) are expressed as mean ± SEM. Statistical analysis was performed using GraphPad Prism 8 (GraphPad Software, San Diego, USA) with either two-tailed unpaired Student’s t test with Welch’s correction, or one-way ANOVA, Dunnett’s test, *P < 0.033, **P < 0.002, ***P < 0.001. Significantly different groups are signified in the respective figure.

## Acknowledgments

The authors are grateful for constructive discussions with Chase Broedersz and Florian Rehfeldt, and for funding by the Deutsche Forschungsgemeinschaft (SFB1032, Projects B02 and B08). Netrin was a kind gift from Raphael Reuten. The authors thank Christian Hohmann from the Nanosystems Initiative Munich (LMU) for the graphical design of Figure 7.

## Figure legends

**Supplementary Fig. S1:**
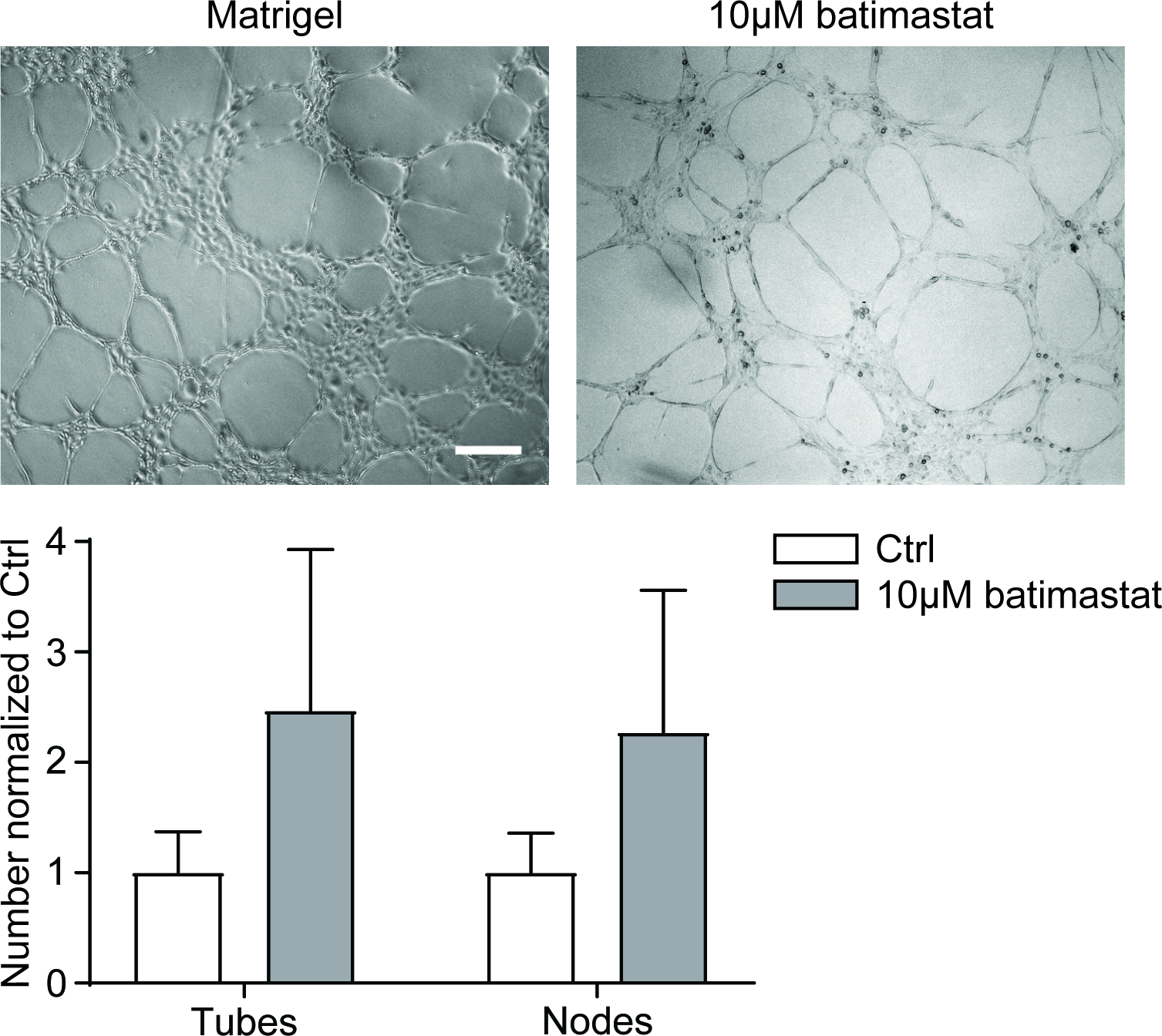
Inhibition of proteolytic activity has no influence on tube formation. HUVECs were seeded on Matrigel and treated with 10 µM batimastat to inhibit a broad range of matrix metalloproteinases (MMPs). Images were taken after 6 h. Quantitative analysis of the number of tubes and nodes normalized to Control (Ctrl) showed that the activity of MMPs is not crucial for HUVEC tube formation. Two-tailed unpaired Student’s t test with Welch’s correction showed no significant differences. Scale bar: 100 µm.

**Supplementary Fig. S2:**
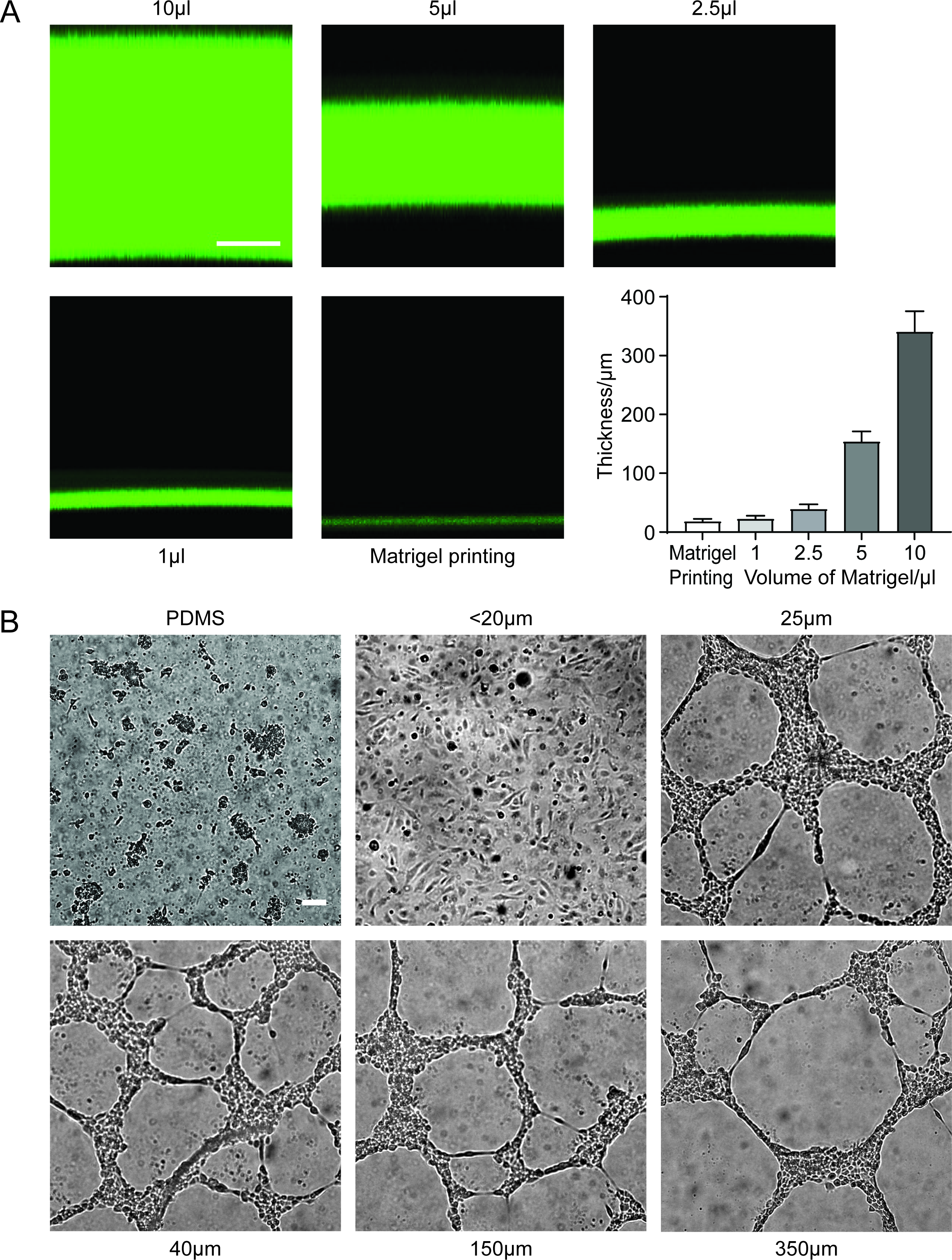
Role of the thickness of the Matrigel layer. **(A)** Matrigel was labelled with a fluorescent dye to measure the gel thickness by confocal microscopy. Different volumes of Matrigel from 1 to 10 µl were added to a PDMS gel, and the thickness was measured. To achieve a thickness of less than 20 µm, Matrigel was printed. Scale bar: 100 µm. **(B)** HUVECs were seeded on a stiff PDMS gel (70 kPa) coated with layers of Matrigel of a different thickness. On uncoated PDMS cells only adhered poorly and formed cell clusters. At a thickness under 20 µm, the cells sensed the stiffness of the underlying PDMS and formed no tubes. In all other cases, the cells did not sense the PDMS and formed the tubular network like on a thick Matrigel layer. Thus, printing of Matrigel was used for all further experiments. Scale bar: 100 µm.

**Supplementary Fig. S3:**
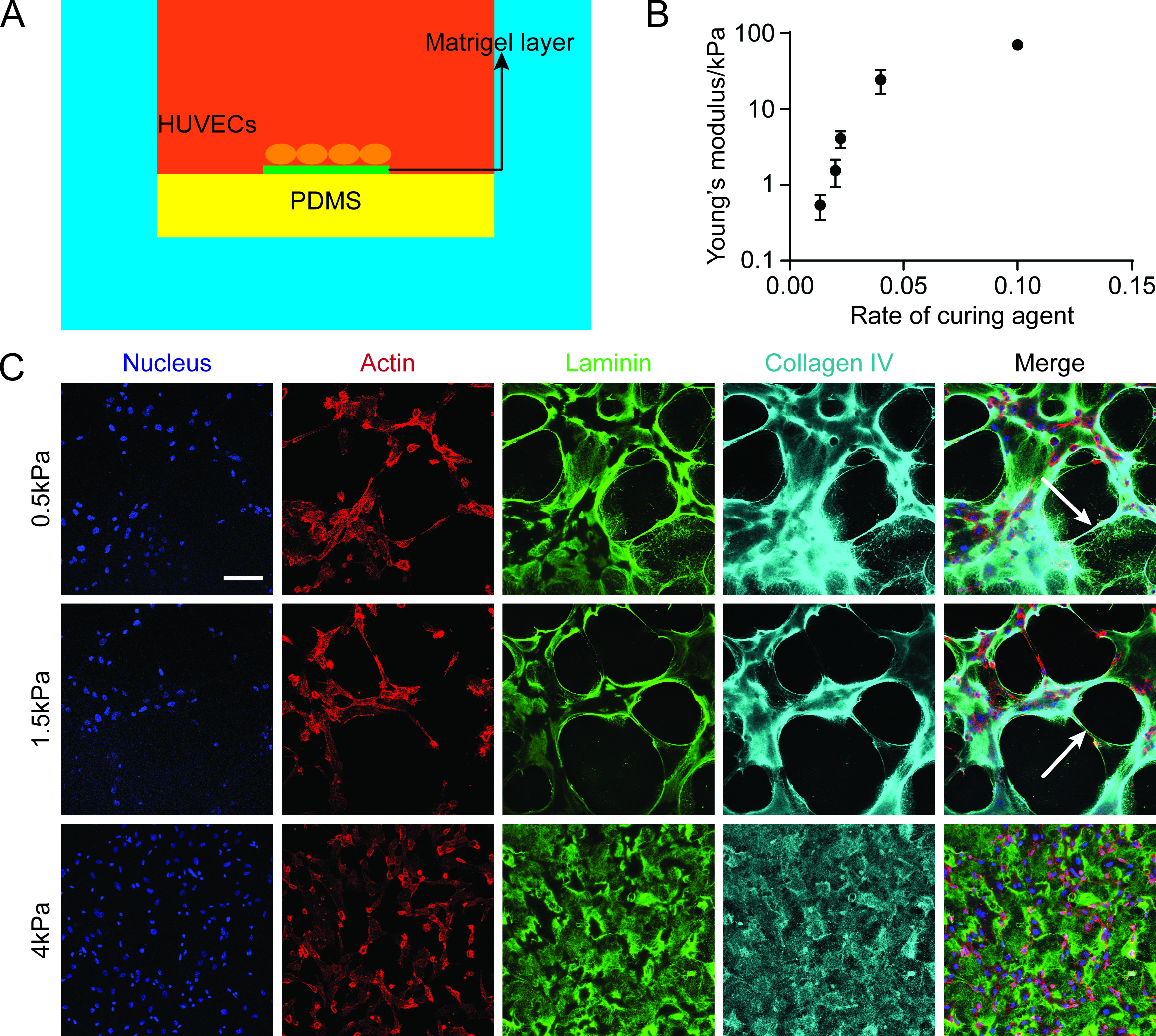
PDMS stiffness influences tube formation. **(A)** Schematic overview of the Matrigel printing on top of a PDMS gel. After activation of the PDMS with the plasma cleaner a thin layer with a thickness under 20 µm was printed on the PDMS. **(B)** PDMS stiffness in dependence of the rate of curing agent. **(C)** Immunostaining of HUVECs seeded on printed Matrigel on top of PDMS gels with different stiffness. If the stiffness was too high (4 kPa), cells did not form a tubular network, but simply spread on the surface. At 0.5 and 1.5 kPa cells interacted and tubes were formed. White arrows indicate the alignment of cells and fibrillary matrix. Scale bar: 100 µm.

**Supplementary Fig. S4:**
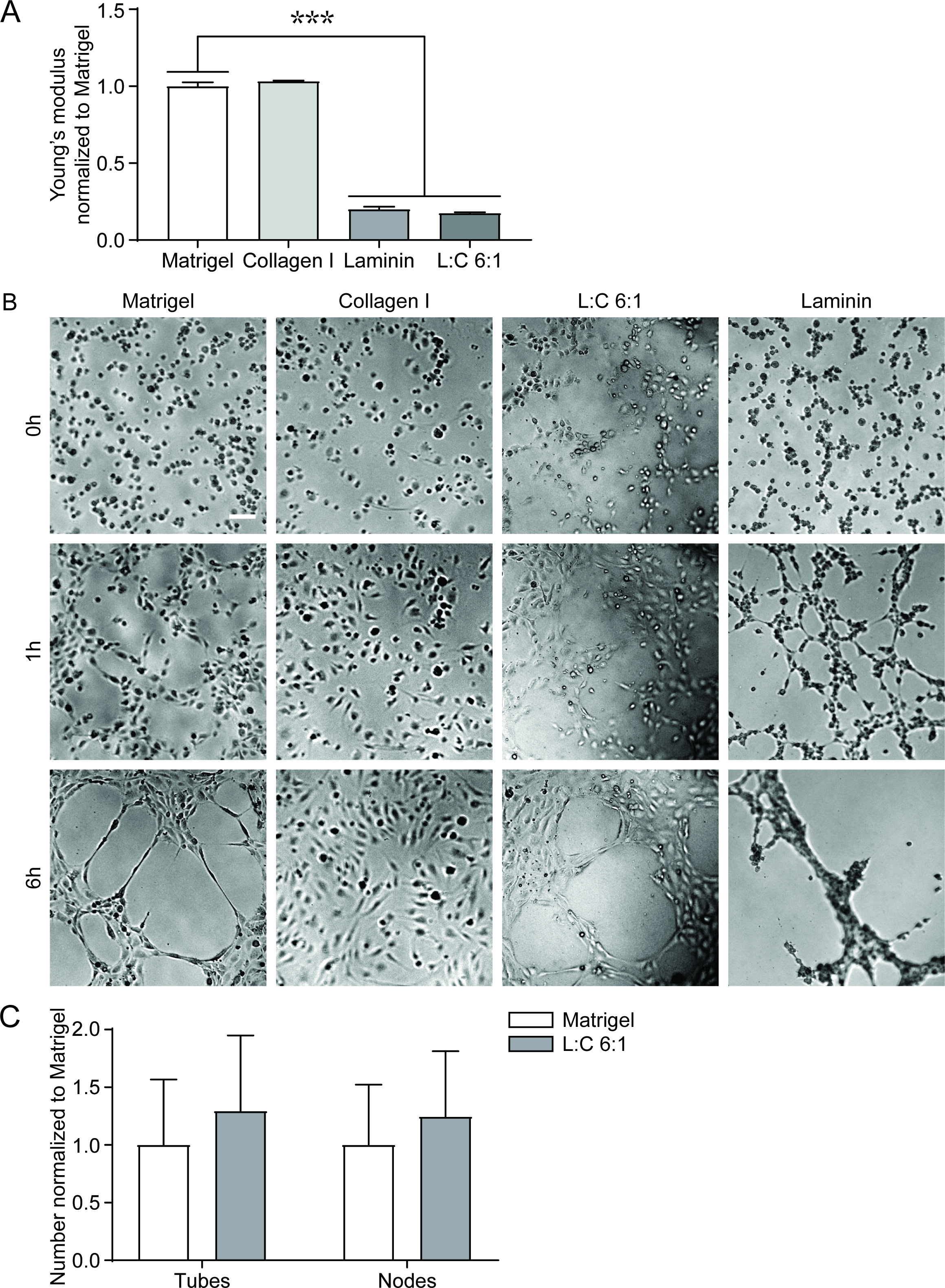
Tube formation on different hydrogel mixtures. **(A)** Young’s modulus of Matrigel, a rat tail collagen I gel (2 mg/ml), a mixture of laminin and collagen I in the ratio 6:1 (L:C 6:1) (4.8 mg/ml:0.8 mg/ml) and a laminin gel (6 mg/ml). The stiffness of Matrigel is significantly higher than the mixture L:C 6:1 or laminin, while there is no difference to collagen. One-way ANOVA, Dunnett’s test, compared to Matrigel, ***P < 0.001. **(B)** HUVECs were seeded on Matrigel, collagen I, L:C 6:1 or laminin. Images were taken after 0 h, 1 h and 6 h. Cells showed the typical tube formation on Matrigel and the L:C 6:1 gel, while on collagen ECs only formed a monolayer. After a short time (1h), the cells on the laminin gel also formed a small network, which then collapsed. Scale bar: 100 µm. **(C)** Quantitative analysis of the number of tubes and nodes normalized to Matrigel showed no difference between the network on Matrigel and the network on L:C 6:1 (Two-tailed unpaired Student’s t test with Welch’s correction).

**Supplementary Fig. S5:**
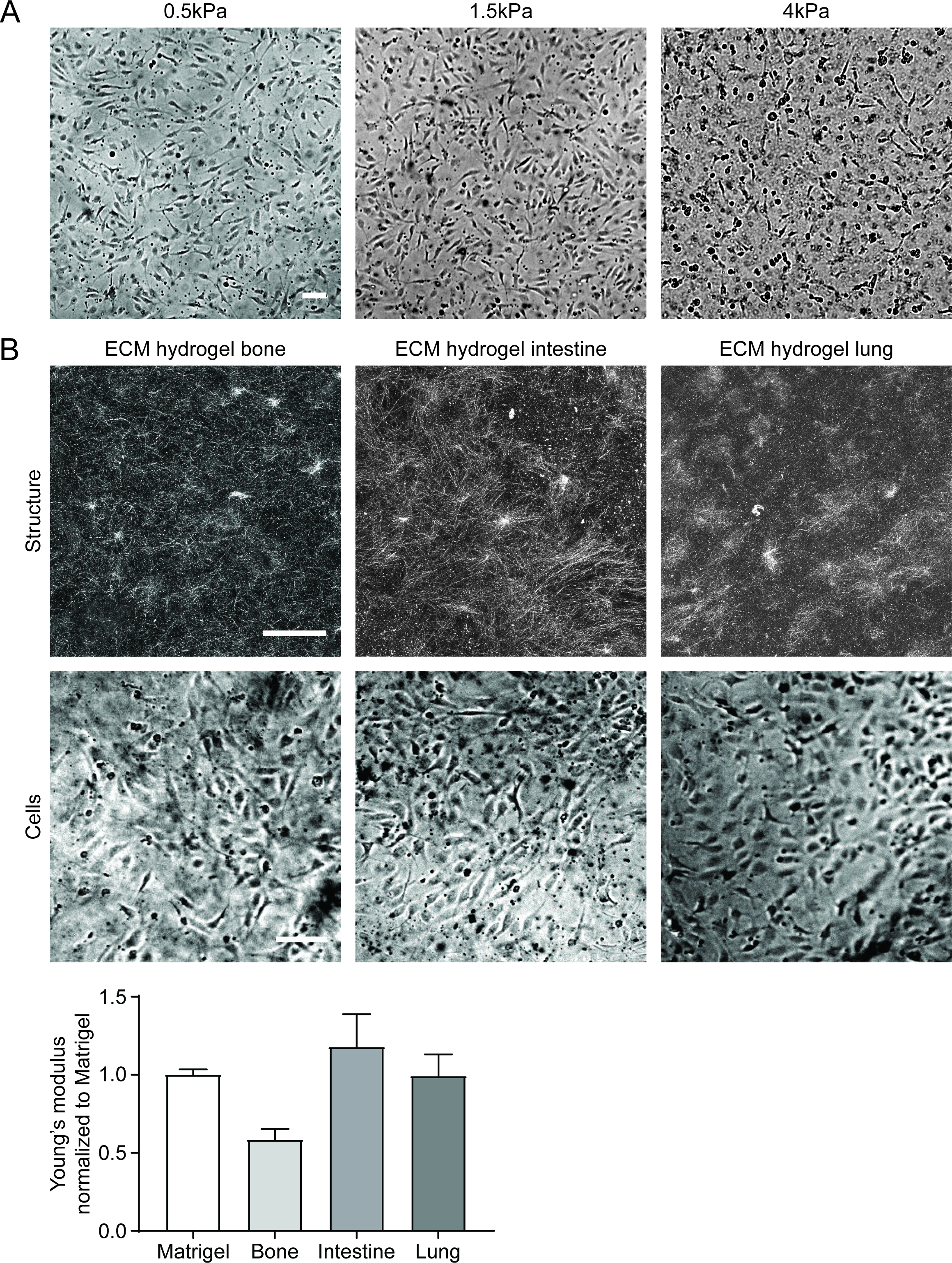
Tube formation on fibrous hydrogels. **(A)** HUVECs were seeded on a collagen I gel printed on top of PDMS gels with different stiffness. In all cases, the cells did not form tubes, but a monolayer. Scale bar: 100 µm. **(B)** First row: total reflection microscopy image of different ECM hydrogel kits from different organs. All hydrogels showed a fibre structure. Scale bar: 100 µm. Second row: Cell behaviour on different ECM hydrogel kits. Cells always formed a monolayer and no tube formation was observed. Scale bar: 100 µm.

**Supplementary Fig. S6:**
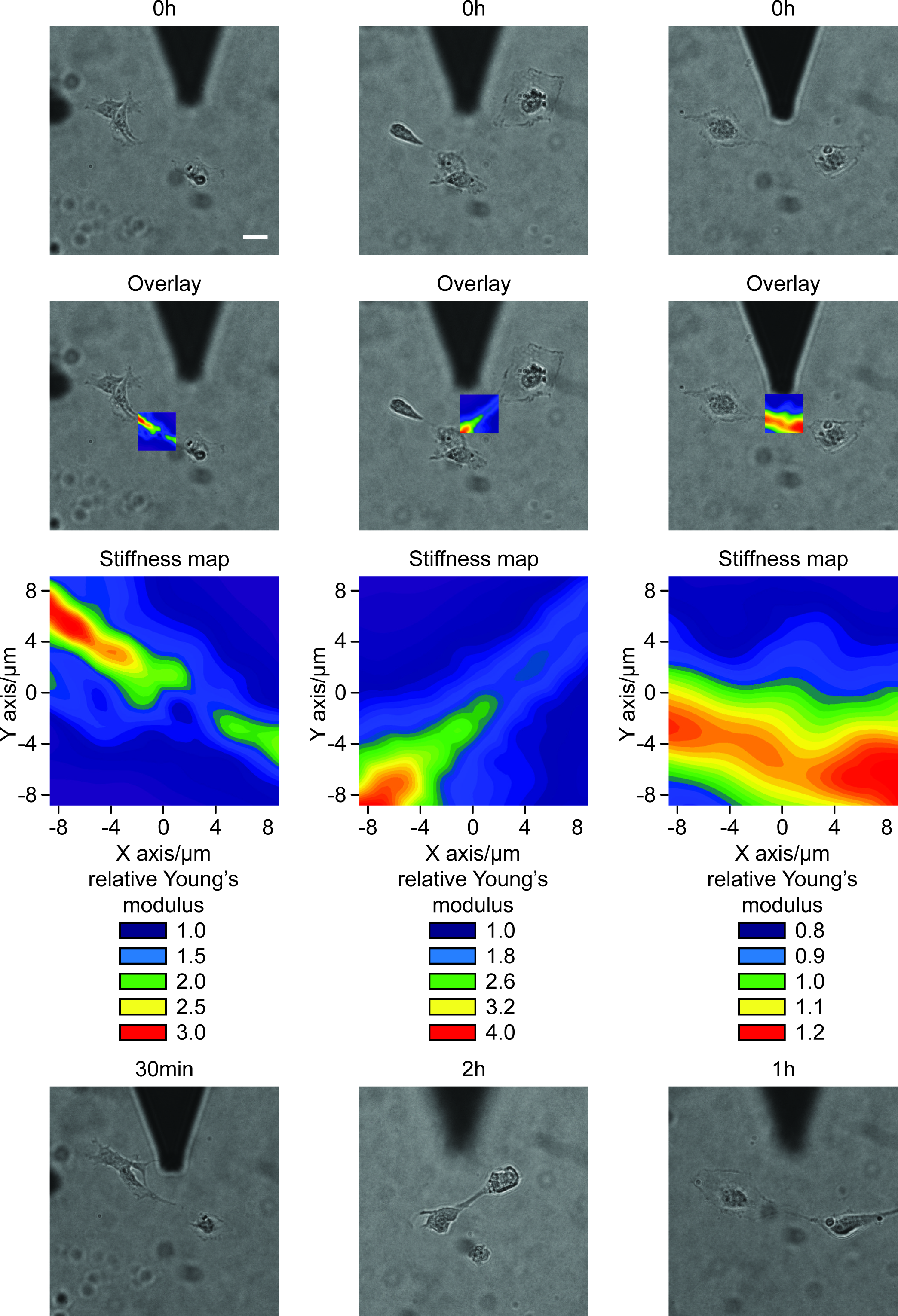
Further examples for the strain stiffening effect. Single cells were seeded on Matrigel and the stiffness of the gel between the cells was measured with an AFM. As control, substrate stiffness in a cell-free area was also measured. The pseudo-coloured stiffness map shows the relative Young’s modulus normalized to the control area. From the overlay of the microscope Image and the stiffness map it can be seen that the cells formed a stiff bridge between them even before cellular protrusions form. In a time range from 30 min to 2 h, the cells used the stiffness line for locating each other and for contacting. Scale bar: 20 µm.

Supplementary Movie S1: Tube formation of endothelial cells on Matrigel.

Supplementary Movie S2: Tubes still form under continuous superfusion.

